# A comprehensive study of mRNA and long noncoding RNAs in Indian Breast cancer patients using transcriptomics approach

**DOI:** 10.1101/2022.04.13.488261

**Authors:** Meghana Manjunath, Snehal Nirgude, Anisha Mhatre, Sai Vemuri Gayatri, Mallika Nataraj, Jayanti Thumsi, Bibha choudhary

**Affiliations:** Institute of Bioinformatics and Applied Biotechnology, Electronic City Phase 1, Bengaluru, Karnataka 560100; BGS Global Hospital, Uttarahalli Main, Bengaluru, Karnataka 560060, India; Manipal Academy of Higher Education, Manipal, Karnataka 576104, India

**Keywords:** Breast Cancer, Transcriptomics, Long-Noncoding RNA, Overall Survival, Gene expression

## Abstract

**Background:** Breast cancer (BC) is one of the leading causes of cancer-associated death in women. Despite the progress in therapeutic regimen, resistance and recurrence of Breast cancer have impacted Overall Survival. Transcriptomic profiling of tumour samples has led to identifying subtype-specific differences, identifying biomarkers, and designing therapeutic strategies. Although there are multiple transcriptomic studies on breast cancer patients from different geographical regions, a comprehensive study on long noncoding RNA (lncRNA) and mRNA in Indian Breast cancer patients in multiple subtypes are very limited. This study aims to understand the subtype-specific alterations and mRNA-lncRNA gene sets.

**Method:** We have performed transcriptome analysis of 17 Indian breast cancer patients and matched normal belonging to 6 different subtypes, i.e., four patients in triple positive, three patients in estrogen receptor-positive (ER+ve), three patients in estrogen and progesterone receptor-positive (ER+ve, PR+ve), two patients in Human epidermal growth factor receptor (Her2+ve), three patients in triple-negative and one patient in ER+ve and Her2 +ve subtypes. Hierarchical clustering and principal component analysis were performed using R packages to derive gene sets. Univariate and multivariate Cox analyses were performed for survival analysis.

**Results:** mRNA and lncRNA expression profiles segregated Indian Breast cancer subtypes with minimum overlap. We have identified a 25mRNA-27 lncRNA gene set, which displayed proper segregation of the subtypes in our data. The same gene set also segregated premenopausal women samples in The Cancer Genome Atlas (TCGA) data. Pathway analysis of the differentially expressed genes revealed unique pathways for premenopausal and postmenopausal women. Kaplan-Meier survival analysis revealed menopausal status, grade of the tumour, and hormonal status displayed statistically significant effects (p < 0.05) on the risk of mortality due to breast cancer. Her2+ve patients showed low overall survival

**Conclusion:** This is the first study describing subtype-specific mRNA and lncRNA gene expression in Indian Breast Cancer patients with unique pathway signatures for premenopausal and postmenopausal breast cancer patients. Additionally, our data identified an mRNA-lncRNA gene set that could segregate pre and postmenopausal women with Breast Cancer. Although the sample size is small, results from this study could be a foundation that could be validated further in a larger dataset to establish an mRNA-lncRNA signature specific to the Indian population which might, in turn, improve therapeutic decisions.

## Background

Breast cancer accounts for 25% of all cancers and exhibits heterogeneity with varied molecular and clinical characteristics (1). The incidence and mortality rate according to GLOBOCAN in 2020, were 34,65,951 new cases and 11,21,413 deaths, respectively worldwide and 1,204,532 new cases and 436,417 deaths in India (2).

Breast cancer is broadly classified based on hormonal status analysis using immunohistochemistry as Luminal A (Progesterone receptor (PR) positive, estrogen receptor (ER) positive and human epidermal growth factor 2 (Her2) negative) and Luminal B (ER-positive, PR positive/negative and Her2 positive) being the estrogen-positive subtypes and Her2 enriched and triple-negative breast cancer (3–5). The fifth subtype is normal-like, resembling normal breast tissue features. Another distinctive subtype showing lower claudin, Epithelial to Mesenchymal markers, and immune receptor expression have been recently identified through molecular analysis (6,7).

Gene expression signatures have been utilized in the past decade for prognosis and to guide treatment in hormone-positive breast cancer patients (8). Oncotype DX, MammaPrint, Prediction Analysis of Microarray 50 (PAM50) are some of the commercially available genomic signatures used in the clinics (9–11). MammaPrint categorizes patients into low and high risk based on the 70-gene profile from microarray (12,13). OncotyeDx is based on the 21 gene expression from the FFPE samples. The relative expression of these genes gives a recurrence score, grouping patients into low risk, intermediate, and high risk (14,15). Prosigna or the PAM50 test depends on the expression of a 50 gene panel that distinguishes the tumour into molecular subtypes and provides the risk of recurrence score (ROR) (16,17). However, these tests have shown success only in caucasian postmenopausal patients and not in younger women with the disease (9,11). Also, these sets have been shown to segregate samples only in microarray data and not in the RNAseq data.

Our understanding of the molecular features of cancer disease has been revolutionized due to the recent advances in next-generation sequencing technology (18), enabling global profiling of mRNAs and non-coding RNAs such as long ncRNAs (lncRNAs), microRNAs, and circRNA. lncRNAs have now been well studied in gene regulation and are known to participate in the development and prognosis of cancer (19–21). Specific mRNA and lncRNA signatures have been associated with different molecular subtypes for breast cancer (22). An Indian cohort study on 543 patients showed that 47% of the BC patients were below 50. 60% of the cohort presented either with HER2+ or TNBC disease (23). Advanced stages of the disease, 51% and 45% of stage III, stage IV belonged to the HER2+ subtype. Recurrence was most frequently observed in HER2+ and TNBC (23). In the present study, survival analysis coupled with Cox has been performed to find the prognostic markers. Kaplan Meier’s Log Rank test and Cox Proportional Hazard regression is a powerful and widely used survival analysis approach (24,25). The molecular heterogeneity of the Indian cohort has not been explored in the subtypes of Breast cancer. This study aims to identify signatures that can stratify BC patients and guide therapy based on altered pathways. Further, identify lncRNA–mRNA regulatory pairs and analyze the probable mechanism of lncRNA involvement in breast cancer progression using in silico tools.

## Methodology

### Study Cohort and Sample classification

The Breast cancer patient samples used for the study were procured from BGS Global hospital, Bengaluru, Karnataka, India. The tumour tissue (n=16) and their respective matched normal (n=17) samples were collected in RNA later. Trizol was added to the samples and stored at -80 until further processing. The samples obtained for the study were histologically classified as Invasive ductal carcinoma (IDC) (except for 1 sample, which was mucinous). The 17 Breast cancer patient samples obtained could be classified into 6 different subtypes based on the expression of estrogen, progesterone, and Her2, which are summarised in Table 1. 10 samples and matched normal samples were also used as a validation cohort. The study was performed under ethical approval from BGS Global Hospitals and IBAB (IEC/Approval/2018-05/06/01A)

**Table 1:** Table depicting sample details of Indian Breast Cancer patients. Odd numbers are matched normals and even numbers are tumour samples. There are a total six subtypes (ER, EH, EP, EPH, Hmod and TNBC) classified based on the expression of estrogen receptor (ER), progesterone receptor (PR) and epidermal growth factor receptor (Her2). IDC (Invasive Ductal Carcinoma)

| Patient_id | Age | Menopause_status | ER | PR | HER2 | Subtype_abbreviation | Type | Recurrence |
| --- | --- | --- | --- | --- | --- | --- | --- | --- |
| P1,2 | 56 | Pre | Positive | Positive | Positive | EPH | IDC | No |
| P3,4 | 56 | Pre | Positive | Positive | Positive | EPH | Mucinous | No |
| P5,6 | 41 | Pre | Negative | Negative | Negative | TNBC | IDC | No |
| P7,8 | 41 | Pre | Positive | Positive | Negative | EP | IDC | No |
| P9,10 | 62 | Post | Positive | Negative | Negative | ER | IDC | No |
| P11,12 | 38 | Pre | Positive | Negative | Negative | ER | IDC | No |
| P13,14 | 48 | Pre | Positive | Negative | Positive | EH | IDC | No |
| P15,16 | 48 | Pre | Positive | Negative | Positive | EH | IDC | Yes |
| P17,18 | 29 | Pre | Negative | Negative | Negative | TNBC | IDC | No |
| P21,22 | 50 | Post | Negative | Negative | Positive | Hmod | IDC | No |
| P23,24 | 35 | Pre | Positive | Positive | Positive | EPH | IDC | Yes |
| P25,26 | 60 | Post | Negative | Negative | Positive | Hmod | IDC | No |
| P27,28 | 60 | Post | Negative | Negative | Negative | TNBC | IDC | No |
| P29,30 | 58 | Post | Positive | Negative | Negative | ER | IDC | No |
| P42T | 60 | Post | Positive | Positive | Negative | EP | IDC | No |
| P43NT | 65 | Post | Positive | Positive | Negative | EP | IDC | No |
| P44NT | 72 | Post | Positive | Positive | Positive | EPH | IDC | No |

### RNA isolation and Library Preparation

Total RNA was extracted using the standard Trizol method from matched tumour and normal samples. RNA was quantitated using QUBIT, and quality was checked using Tapestation. mRNA libraries were prepared using Illumina TruSeq RNA Library Prep Kit v2. Briefly, mRNA was isolated using oligo-dT beads and followed by fragmentation. Fragmented RNA was then converted to cDNA, and adaptor ligation was performed. Size selection was performed on Adaptor ligated libraries using ampure beads. The libraries were then amplified and checked on a tape station to determine the library size.

### RNA sequencing and Data analysis

The samples were sequenced in-house using Illumina Hiseq2500 to acquire100bp paired-end reads. Samples had reads >10 million (Supplementary Table 1). The quality of the reads was checked using the Fastqc tool (26). The reads were then aligned to the reference hg38 (Downloaded from The University of California, Santa Cruz (UCSC) genome browser) using bowtie2 with default parameters (27). A SAM (Sequence alignment map) format file was obtained as an output of the bowtie2. A binary alignment map (BAM) file was obtained using Samtools (28) from the SAM file. hg38refseq.bed annotation file was downloaded from UCSC and read counts were generated using bed tools (29). The read counts for each matched normal and tumour pair were given as an input to DESeq, an R package to obtain differentially expressed genes (30).

### Pathway enrichment analysis

A cutoff of p-value less than 0.05 and log2 foldchange (<-1 and >+1) was used to obtain a significant DEG list for each normal tumour sample pair. Common significant DEGs from patients belonging to each subtype were taken out and subjected to the Reactome pathway analysis (https://reactome.org/) to obtain subtype-specific upregulated and downregulated signature pathways. Also, Pre-menopause and post-menopause signature pathways. Pathways with a False discovery rate of less than 0.1 have been plotted in a bubble plot using ggplot2, an R package.

### LncRNA analysis

The Bam files obtained for each tumour and their respective matched normal samples from samtools were given as an input to bedtools with gencode.v34.long_noncoding_RNAs.gtf annotation file obtained from GENCODE (https://www.gencodegenes.org/human/release_34.html). The read count file was for each tumour-normal pair was given as an input to DESeq (R package) to obtain differentially expressed LncRNAs. LncRNAs were then compared against the lnc2cancer database (31) and the known breast cancer-related lncRNAs were selected.

To obtain LncRNA-mRNA pairs, for each subtype, the list of unique lncRNAs with genomic regions information and NCBI RefSeq hg38 reference was given as an input to bed tools intersect. Bedtools intersect is useful to screen for overlaps between two sets of genomic features. To generate potential overlapping (anti-sense) lncRNA-mRNA pairs, a window greater than 1000 bases were selected.

### Sample distance calculation

‘dist’ in R was used to calculate the Euclidean distance between samples. Principal Component Analysis (PCA) was performed on all patient samples using Deseq2’s plot PCA function with different subtypes as the variable of interest. Significant Genes from different patient samples with differentially expressed (DE) genes were sorted based on p-value and log2Fold change. Heatmap was plotted for the filtered genes using pheatmap with default Euclidean distance parameters. To determine the overall similarity and signature of breast cancer patient subtypes gene expression profiles, hierarchical clustering was performed between all samples and was visualized using the pheatmap.

### Survival analysis

Clinical parameters of 381 Indian breast cancer patients were obtained from BGS Global hospital. Out of which 17 samples were sequenced. To investigate the impact of the clinical parameters such as menopausal status, age, stage, and grade of tumour and therapy on the prognostic survival of breast cancer patients, KM survival curve analysis was performed, and Cox proportional hazard ratio (HR with 95% CI) plots. The calculations were done using univariate and multivariate Cox analysis with Survminer (https://github.com/kassambara/survminer) and survival packages of R (24).

### Extraction of Breast cancer expression data from TCGA

RSEM values for 946 Breast cancer patients file (BRCA.rnaseqv2_illuminahiseq_rnaseqv2_unc_edu Level_3_RSEM_genes_normalized_data. data.txt) were downloaded from TCGA (http://firebrowse.org/?cohort=BRCA#). The file having barcode information for each patient was also procured from TCGA (https://portal.gdc.cancer.gov/) to obtain hormone receptor subtype information.

### First-strand cDNA synthesis

Once the intact RNA was obtained, complementary DNA (cDNA) synthesis was initiated. For synthesizing cDNA from mRNA random hexamers were used. For making cDNA, 4 μg of RNA was taken from each patient sample from the validation cohort. To remove DNA contamination, the RNA samples were treated with DNases (37°C, 10 min) and proceeded to cDNA synthesis using M-MuLV reverse transcriptase (37°C, 1h). Initially, RNA samples were incubated with adaptor primers and dNTPs for 1 hr at 37°C. Followed by the addition of random hexamers and incubated for 1 hr at 37°C (32). For negative control, a reaction without RTase (reverse transcriptase) was kept for each sample.

### Real-time PCR for investigating the expression of marker genes

Real-time PCR was conducted using SYBR^®^ Green chemistry (33). Primers for BCL2, BRCA1, TP53, CD44l, CD44s, ALDH1A and HOTAIR genes were used with GAPDH primer as an internal control. The initial denaturation was done at 95°C for 5 minutes and followed by the cycling stage (40 cycles, 95°C for 20 sec, 53°C for 20 sec, 72°C for 20 sec) and melt curve stage (34).

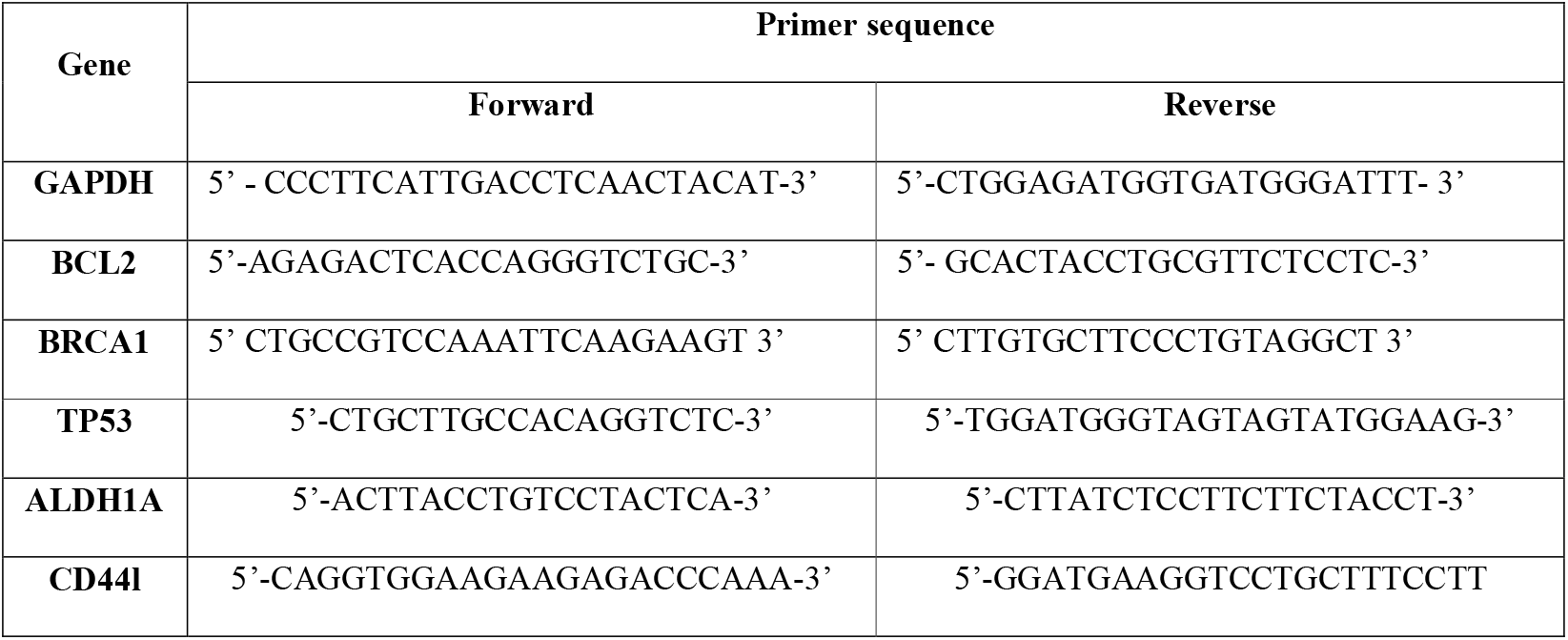
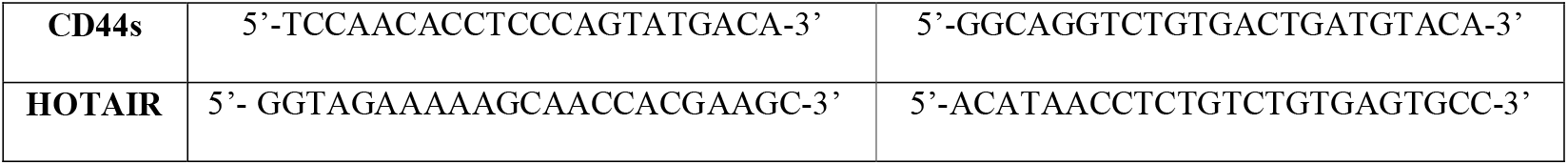

Here the relative gene expression is calculated by correlating the expression of the housekeeping gene and the expression of the target gene in the control/normal sample. Ct is the cycle number at which the fluorescence crosses the threshold level (35,36). The equation for relative quantitation (RQ) value is:

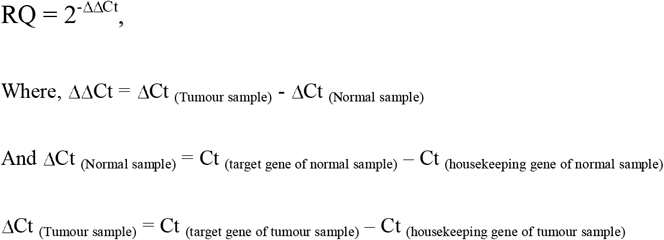

Graphs showing relative quantification for all the samples were plotted using the GraphPad prism software (37).

### Statistical analysis

Statistical analyses and graphing were done using GraphPad Prism 7.0 software (GraphPad, San Diego, CA, USA) and R packages. Deseq2 uses Wald test statistic with a probability to generate a significant gene list. The Benjamini–Hochberg False Discovery Rate (FDR) method was used for choosing significant pathways from the Reactome database. For comparative qRT-PCR analysis, a two-tailed t-test was applied to calculate the significance. If the p-value was less than 0.05, the results were significant.

## Results

### Her2 positive patients and disease recurrent subgroup had poor survival among the Breast cancer subtypes in the Indian cohort

Kaplan-Meier plots were used to depict survival for different clinical parameters of Indian Breast cancer patients. Properties like menopausal status, hormone receptor status, tumour grade, recurrence and stages were analysed among the cohort. There was a total of 381 patients with data available for menopause status and among them, there were two groups: pre (n=216) and post (n=159). Significantly (p=0.11) low survival was observed for premenopausal patients compared to postmenopausal samples (Figure 1a). A multivariate Cox proportional hazards analysis of the menopause status revealed that postmenopausal patients displayed a hazard ratio of 0.51 indicating that this group of patients has a high risk of death (Figure 1b). When the disease recurrence parameter was checked, the recurrent patients were divided into various categories like local, distant, distant + regional, local + distant + regionally based on where the recurrence has occurred. When all these categories were compared, local recurrence had poor survival (p-value< 0.0001) followed by distant recurrence and local + distant + regional group (Figure 1c). Breast cancer is classified commonly based on the expression of hormone receptors. Within the hormone receptors subtypes, Her2 positive subtype had worse prognoses (p=0.026<0.05) (Figure 1e). A multivariate Cox proportional hazards analysis of the hormone receptor subtypes pointed out that Her2 positive patients displayed a significant (p-value 0.097) hazard ratio of 2.4 indicating that this subtype has a high risk of death (Figure 1d). Among the different stages in our cohort, it was observed that stage IV exhibited worse survival in comparison to others with a significance of p-value< 0.0001 (Figure 1f). For analysing survival of tumour grade, patients falling in grades 1, 2 and 3 were plotted. Patients with higher-grade had low survival in comparison to grades 1 and 2 (p-value-0.011) (Figure 1g). Among the 381samples used for analysis, 17 matched tumour-normal samples were subjected to RNA sequencing analysis to identify differentially expressed genes and pathways regulated in the presence/absence of hormone, pre and postmenopausal samples irrespective of hormone status

**Figure 1:**
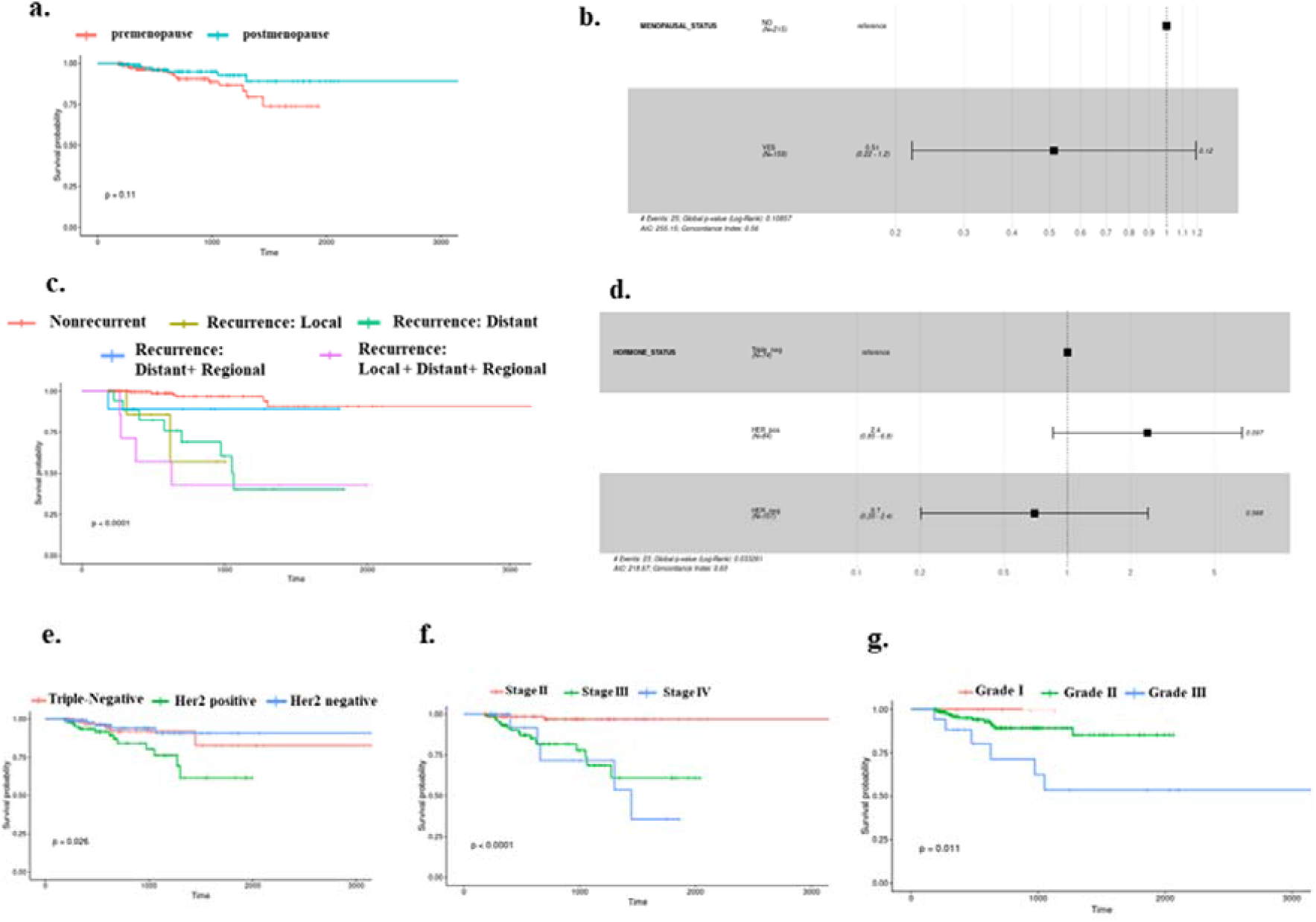
Kaplan-Meier survival plots showing differences in probabilities between various clinical parameters. **a**. This plot depicts a low survival for postmenopausal samples compared to premenopausal women samples **b**. A cox-proportional hazard ratios plot, showing significant variations between pre and postmenopausal status **c**. This survival plot shows differential probabilities between different types of recurrent and non-recurrent samples. **d**. A cox-proportional hazard ratios plot, showing significant variations between Triple Negative, Her2 positive and Her2 negative subtypes of Breast Cancer **e**. Survival plot for Triple Negative, Her2 positive and Her2 negative subtypes of Breast Cancer. **f**. Survival plot for different stages of Breast cancer. **g**. The plot is for displaying different survival probabilities for samples belonging to different tumour grade.

### Gene expression and unique pathways alterations segregate six breast cancer subtypes

Differential gene expression analysis was performed on tumour and matched-normal ER (3pairs), EP (3 pairs), EPH (4 pairs), Hmod (2 pairs), EH (2pairs), and TNBC (3pairs) patients. Among 6 subtypes, ER subtype had a maximum alteration in gene expression where 2572 genes were uniquely significantly downregulated and 1324 upregulated (log2fold change ≤ and ≥□□1) followed by EP (543 down and 795up), Hmod (514, 373), EPH (183 and 243), TNBC (116 and 173), and least in EH (31 and 37) (Figure 2a). As expected, a minimal overlap was observed between the subtypes, with ER having maximum overlap with EPH, EP, and EH (Figure 2b). It is well known that a balance tilt in oncogenic (ONC)/tumour suppressor (TSG) drives oncogenesis; we checked for the alteration in ONC and TS across the subtypes. The DE genes were subjected to oncogene/tumour suppressor analysis using breast cancer-specific oncogenes (https://oncovar.org/) and tumour suppressors (https://bioinfo.uth.edu/TSGene/). Each of the subtypes was analyzed for upregulated oncogenes and downregulated tumour suppressors. Most downregulated TSGs, and upregulated oncogenes were observed in the EH (16%TSG and 27%ONC), followed by TNBC (7.7%TSG and 5.6%ONC). The fewest alterations were observed in Hmod (0.97%TSG and 1.3% ONC), while EPH (4.37% TSG and 4.9% ONC), EP (6.9%TSG and 2.7%ONC), and ER (2.7%TSG and 5%ONC), indicating differences in the alterations in Oncogenes and tumour suppressors among breast cancer subtypes (Figure 2c). Figure 2d depicts the list of significantly upregulated oncogenes and downregulated tumour suppressor genes in each subtype. Oncogenes like MYC, SIRT6, IL7R, CCNE1, PAX8, BCL11A were upregulated and TSGs DUSP1, AGTR1, NOTCH2, CREBBP, ITGA7 were downregulated in the subtypes. Further, to identify the deregulated pathways; Upregulated and downregulated genes for each subtype were given as an input separately to the Reactome database, and the results were filtered for p-value < 0.01, and the pathways with a gene count of more than 3 were selected. The top results are represented in a bubble plot. Among the notably affected pathways are downregulated Keratinisation and RUNX3 related pathways among the ER samples; downregulated ubiquitination and upregulated FGFR signalling among Hmod; ECM interactions and notch signalling downregulated in TNBC and upregulated collagen and cellular pathways. AP2 related genes are regulated in opposite directions in ER and Hmod (Supplementary Figures 1 and 2). The difference observed in survival between pre and postmenopausal and the understanding that premenopausal disease is aggressive, we identified pathways that regulate these phenotypes.

### Pre and postmenopausal samples show unique pathway signatures

The Breast cancer patient samples were divided into 2 categories, pre and postmenopausal, based on the menopause data from the clinical features procured from the hospital. Genes with log2foldchange < 1 and >-1 were filtered out for each patient. Common DEGs were pulled out from patients from each group and then further analysed. Venn was performed to identify common and unique genes among the two types (Supplementary File 1). Pre and post groups showed unique 72 and 380 downregulated and 71 and 311 unique upregulated genes respectively than common genes (1 and 2 genes for down and upregulated respectively) (Figure 2f). These unique genes were then checked for chromosome distribution and found that downregulated genes were on chromosomes 5, 17, and 2 in the post and premenopausal samples, respectively (Figure 2e). Upregulated genes were primarily present in chromosome 1 for both post and premenopausal samples, with chromosome 12 being additional for the post. These unique significant upregulated and downregulated genes were given as an input separately to the Reactome database to obtain deregulated pathways. In postmenopausal women, Breast cancer samples, pathways related to metabolism like phospholipid metabolism, amino acid metabolism, glycogen metabolism were downregulated, and Cell cycle processes connected to transcription, translation were upregulated. In the case of premenopausal women Breast cancer samples, extracellular matrix regulation, collagen dependent pathways were downregulated, and single and double-stranded DNA repair pathways and immune-related pathways were upregulated (Figure 2g). To understand the gene signatures signifying each subtype also correlated with already existing molecular panels such as PAM50, Oncotypedx, and MammaPrint, we performed PCA of the samples with select gene sets. Pre and postmenopausal samples show unique pathway signatures.

**Figure 2:**
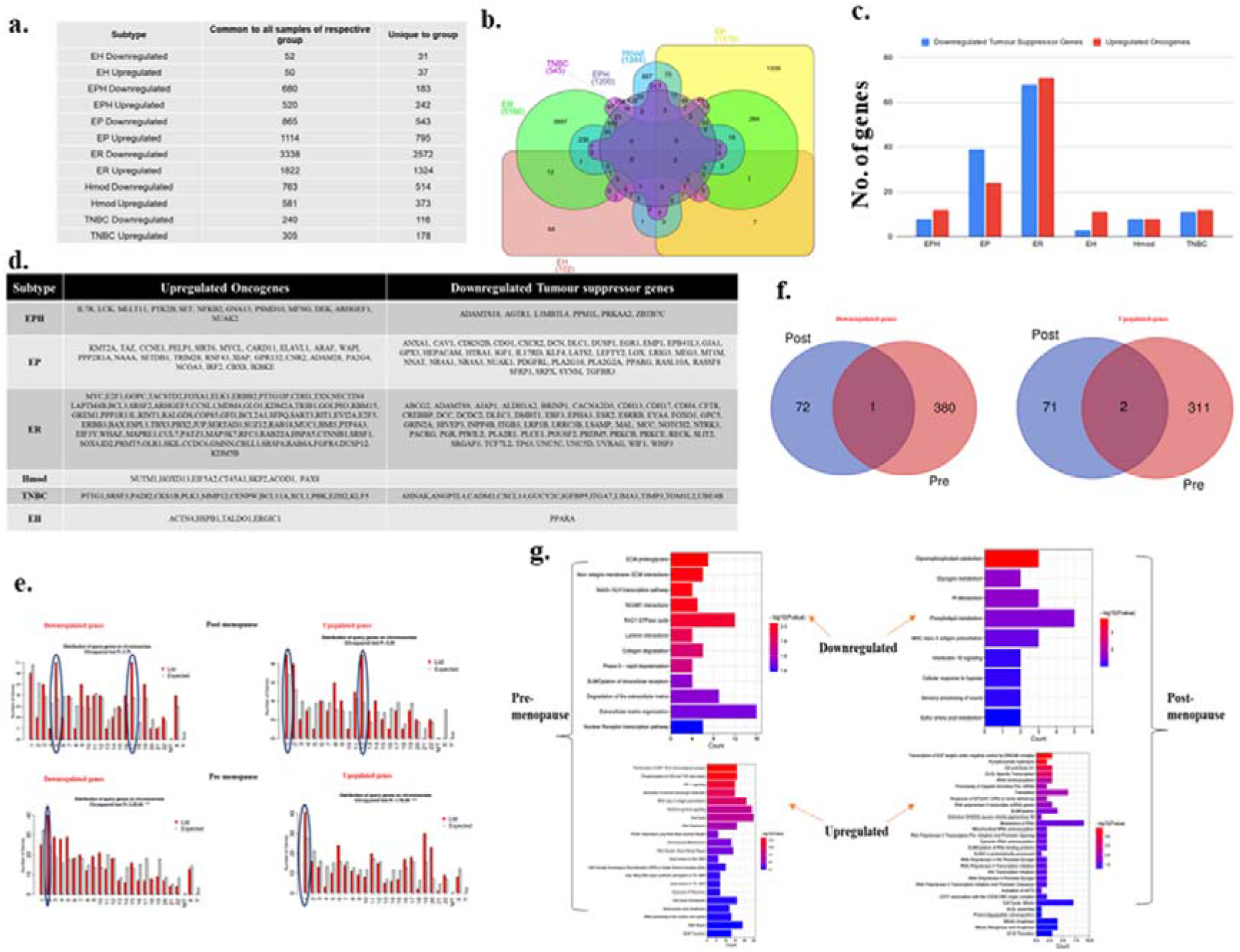
**a.** Table showing the number of differentially expressed genes that are common to all patients in a subtype and unique genes compared to other subtypes of breast cancer **b**. Venn diagram showing common and unique genes among different subtypes of breast cancer patients **c**. A bar graph depicting No. of upregulated oncogenes and downregulated tumour suppressor genes in six subtypes of Indian Breast cancer patient samples. d. A table with a list of upregulated oncogenes and downregulated tumour suppressor genes **e**.Venn diagram showing common and unique genes among pre-menopause and post-menopause Indian Breast cancer patient samples. **f**. Bar graphs depicting gene distribution on chromosomes in pre and post-menopause Indian Breast cancer patient samples. **g**. Bar graphs representing significantly upregulated and downregulated pathways in pre and post-menopause Indian Breast cancer patient samples. Y-axis shows pathway terms and the x-axis is gene count. The colour gradient of the bar is based on the p-value.

### A 25 gene set was identified for the Indian Breast Cancer cohort

Although gene expression patterns were unique for each subtype, when Principal component analysis was performed for all the genes in each sample, no segregation was observed. We performed PCA PAM50, MammaPrint, and OncotypeDX gene sets using RNA-seq data of the Indian cohort. No clear segregation of the subtypes was observed in the Indian cohort, suggesting differences in microarray and RNA-Deseq based analysis (Figure 3 a-c). To check if the absence of segregation can be due to differences in the technology used, we downloaded RNAseq data from TCGA and analyzed for these gene sets; clear segregation was observed among hormone positive and negative samples (Supplementary Figure 3 a-c) in the PCA, suggesting that it is not dependent on the technology used. Samples from TCGA mainly belong to the Caucasian population, showing a distinct separation. The panels are designed primarily for a specific population, suggesting population-specific expressions that may underlie the differences observed.

**Figure 3:**
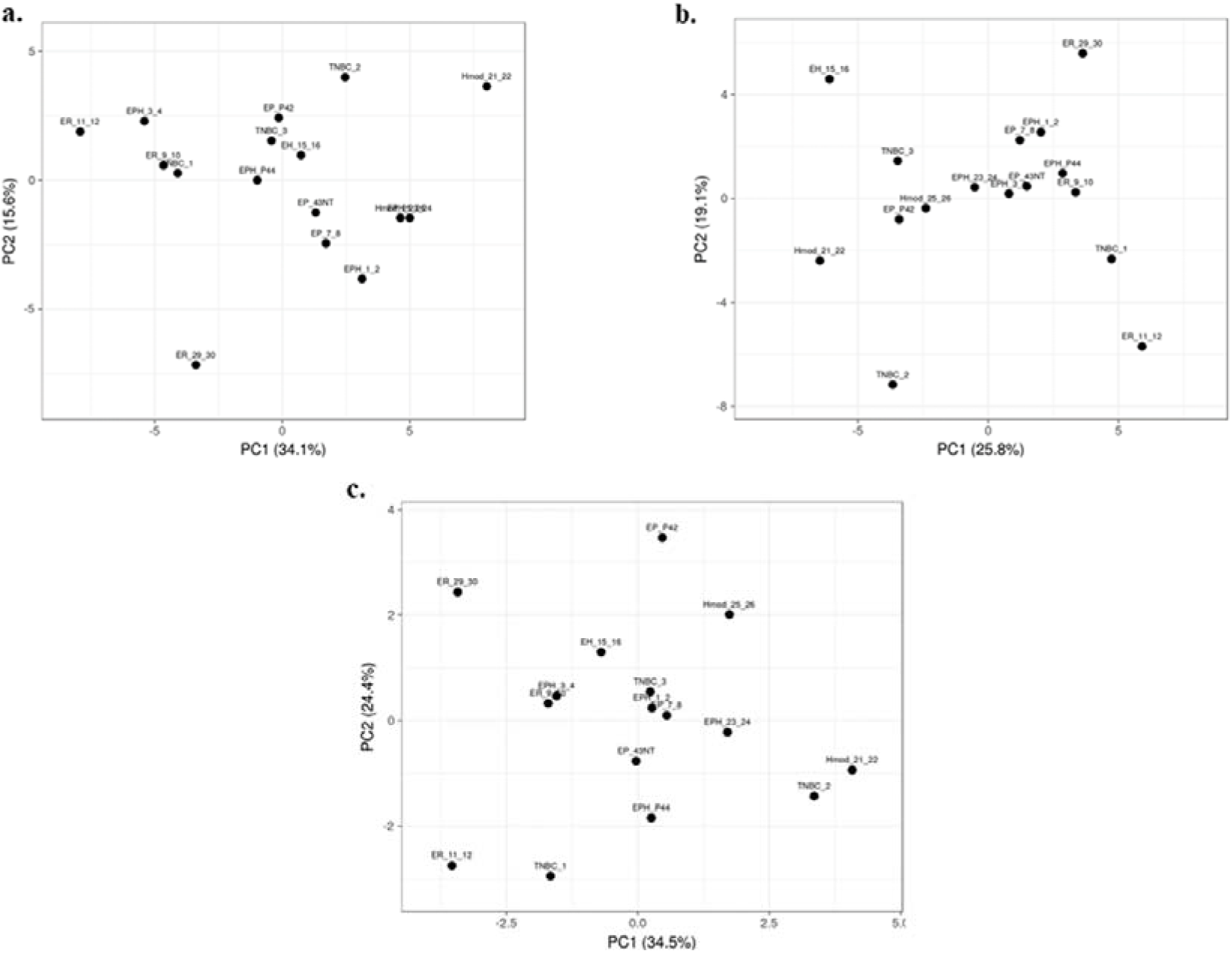
Principal Component Analysis of Indian breast cancer patient samples with the **a**. PAM50 **b**. MammaPrint and **c**. OncotypeDX gene sets.

To narrow down to a gene set that might segregate the subtypes, the list of genes was narrowed down based on log2fold change, p-value, and a significant DEG list for each patient was obtained. Significant DEGs common to all the patients belonging to each subtype were taken out. These common DEGs were then compared between the subtypes, and a unique set of genes was procured for each of the subtypes. Among the unique DEG list, genes already known in the existing literature relevant to cancer were narrowed down. Principal Component Analysis and heatmaps were iteratively used to narrow these lists into combinations that segregated the patient samples into their distinct hormone receptor-based subtypes. 25 mRNAs were identified specific to our data (Figure 4 a, b). The selected mRNAs show proper segregation in the PCA of hormone subtypes (Figure 4 c). Also, the candidate gene set was used to evaluate segregation between pre and postmenopausal women samples in the Indian cohort. Postmenopausal breast cancer samples showed better segregation in the PCA of the subtypes than premenopausal samples (Figure 4d). This gene set was then checked in TCGA samples. RSEM normalized values for 946 individuals were downloaded from TCGA. The samples were segregated into pre and postmenopausal. PCA plot for a 25 gene set was plotted for TCGA samples and observed that premenopausal samples showed better segregation than postmenopausal samples. The existing gene sets could only segregate postmenopausal TCGA samples. Hence, the 25 gene set could be used for premenopausal women in the Caucasian population (Supplementary Figure 3d). Additionally, genes such as CNR2 (Luminal B), LRRC3B, EYA4, TMEFF2 (Luminal A), ESR2, GRIN2A, ERBB4, NNAT (ER-ve a.k.a Hmod and TNBC) also differ significantly between TCGA and Indian data. These are therefore of particular interest as population-specific markers. Since we did not find segregation of the premenopausal samples and understand if the addition of lncRNA to the panel improves segregation of the breast cancer subtypes, we performed differential lncRNA analysis across subtypes.

**Figure 4:**
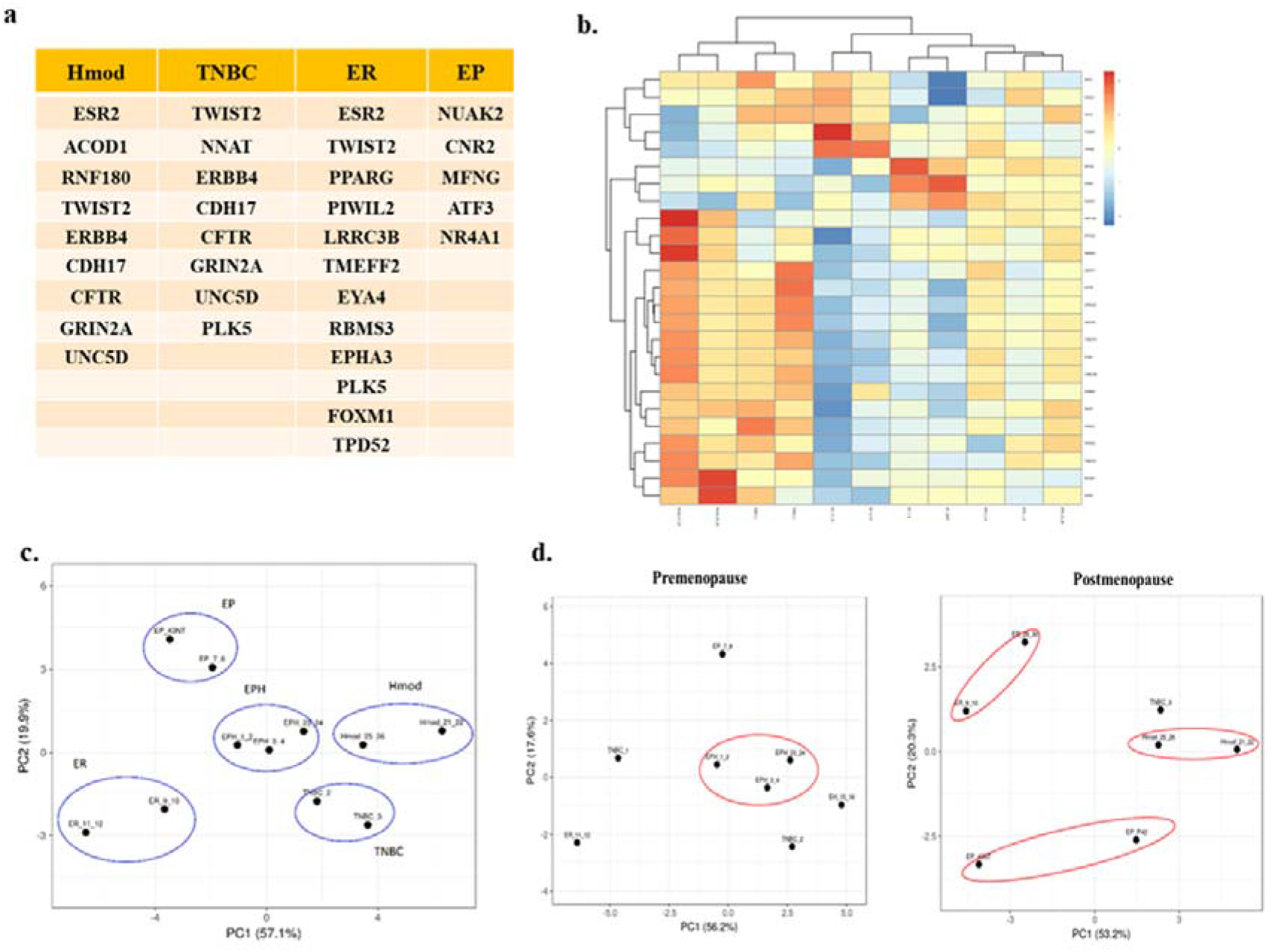
**a.** Table depicting unique mRNAs with potential as Indian-specific biomarkers derived from different subtypes. **b**. Heatmap of unique mRNAs chosen as potential biomarkers in the Indian population. Blue colour represents downregulated genes and red represents upregulation. **c**. PCA plot showing segregation of Indian patients with selected mRNAs. **d**. PCA plots showing segregation with selected mRNAs between pre and post-menopause Indian Breast cancer patients.

### Unique lncRNA expression pattern in Indian Breast cancer subtypes

lncRNA is known to regulate gene expression and is known for its tissue-specific expression (38,39). To identify subtype-specific lncRNA, DESeq was performed using matched normal/ tumour pairs for each sample and obtained lncRNAs, which were either upregulated/ downregulated in all samples of a group. Among the subtypes, ER showed the most significant number of alterations in lncRNA as was observed for mRNA, followed by Hmod and EP, and the least in EH. Triple-negative and triple-positive cancer showed comparable levels of alterations in both up-and down-regulated lncRNAs (Figure 5a). When commonly regulated differential lncRNAs were looked into using the Venn diagram, it was found that no lncRNA was common to all subtypes (Figure 5b), indicating subtype specificity of lncRNA. TRG-AS1, MAFA-AS1, MELTF-AS1 in EPH, TET-AS1 ZNF26-DT, C4A-AS1 in TNBC, FZD4-AS1, CHL1-As1, B4GALT1-AS1 in Hood, HOTAIR, EGOT, FOXN3-AS2, TMEM12-AS1 in ER, DOCK9-AS1, MORC1-AS1, GASAL1 in EP were some of the uniquely upregulated lncRNAs in a subtype-specific manner. ARNTl2-AS1, ELMO-AS1, NAMA in EPH, B4GALT1-AS1, HOXB-AS1, EP300-AS1 in TNBC, NCF4-AS1, ZSWIM8-AS1, DICER1-AS1 in Hood, NRIR, TP53TG1, DDX11-AS1 in ER, MYLK-AS1, ADNP-AS1, SNHG12, HNF4A-AS1 in EP (Supplementary File 2) were some of the uniquely downregulated lncRNAs in subtypes.

**Figure 5:**
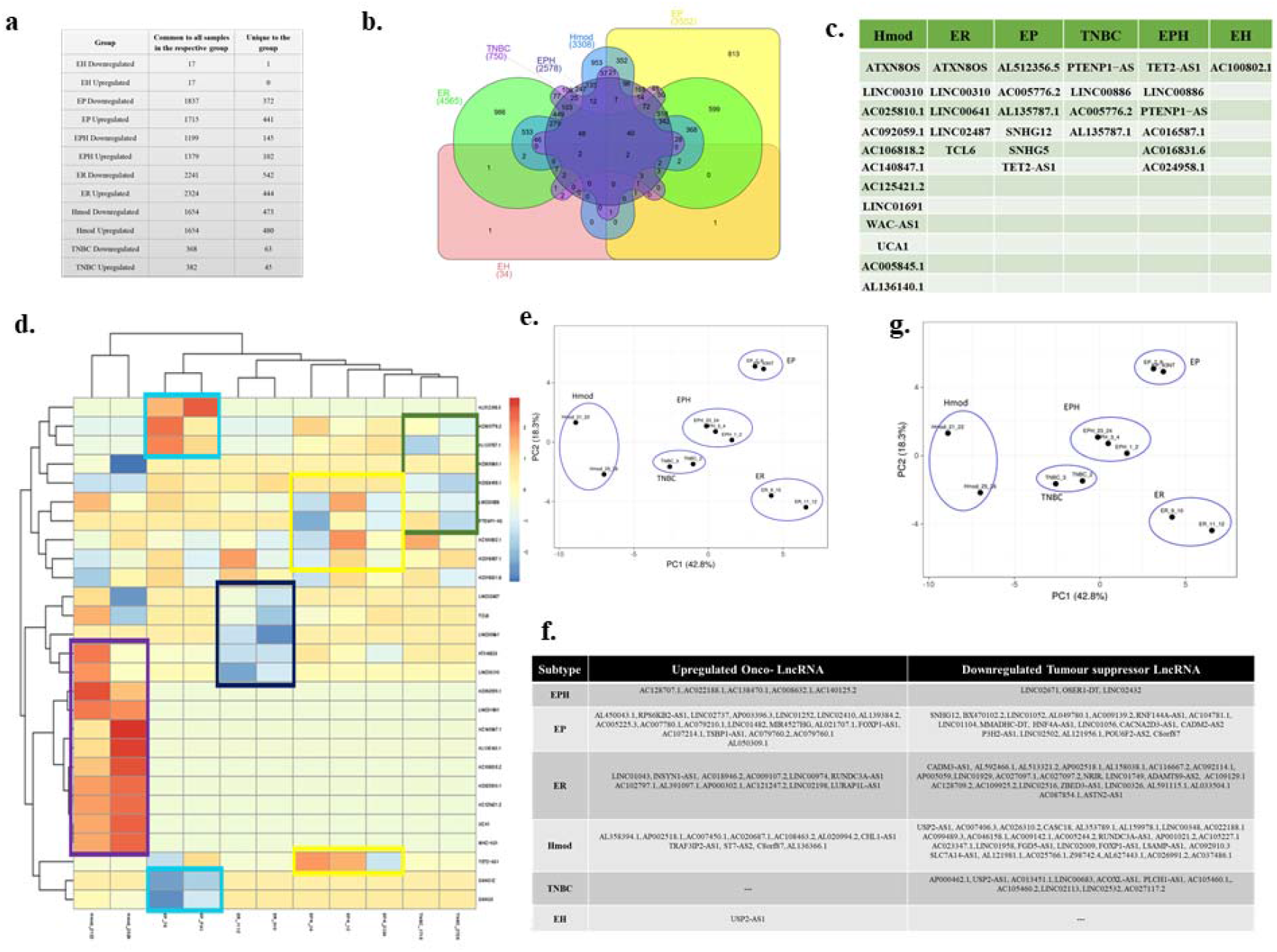
**a.**Table showing the number of differentially expressed LncRNAs that are common to all patients in a subtype and unique LncRNAs compared to other subtypes of breast cancer **b**. Venn diagram showing common and unique LncRNAs among different subtypes of breast cancer patients. **c**. Table depicting unique LncRNAs with potential as Indian-specific biomarkers derived from different subtypes. **d**. Heatmap of unique LncRNAs chosen as potential biomarkers in the Indian population. Blue colour represents downregulated genes and red represents upregulation. **e**. PCA plot showing segregation of Indian patients with selected LncRNAs. **f**. Table showing onco and tumour suppressor lncRNAs segregated subtype wise **g**. PCA plot of combined signature of mRNA and lncRNA for Indian Breast Cancer patient samples.

To identify differentially expressed lncRNA between pre and postmenopausal samples, DE lncRNA was obtained from pre, and postmenopausal samples and common and unique lncRNA were obtained. AL357054.2 was the only LncRNA commonly upregulated among postmenopausal samples. LINC02306, AL442163.1, AC124947.1, and AC016831.1 were commonly downregulated, and AC024958.1, AC011447.3 were commonly upregulated in Premenopausal samples (Supplementary File 1).

Just as in mRNA analysis, the oncogenes and tumour suppressors regulate tumourigenesis; we also classified the lncRNA as ONC and TSG and identified subtype-specific lncRNA. (Figure 5f). The unique lncRNAs were analysed for each subtype and were compared against the lnc2cancer database, and the known breast cancer-related lncRNAs were selected. A set of 27 lncRNA was identified from the data (Figure 5c). This gene set was devised iteratively following the removal of frequent outliers. It was observed that most lncRNA was upregulated in Hmod, whereas the same had negligible expression in all other subtypes. Similarly, ER showed downregulated lncRNA, which was upregulated in other subtypes. The expression pattern using lncRNA showed an apparent demarcation, as shown in the heatmap (Figure 5d). As depicted in Figure 5c, ATXN8OS, UCA1, SNHG12, SNHG5, LINC02487, TCL6, TET2-AS1, PTENP1−AS were the identified lncRNAs from our data. When the same lncRNA set was compared in pre, and postmenopausal women samples, Hmod and ER subtypes segregated better in the post, and premenopausal samples did not show any clear pattern in the PCA (Supplementary Figure 3e). Since lncRNA signatures also segregated the subtypes only in postmenopausal samples, we checked for the segregation using mRNA and lncRNA signatures.

### 25mRNA and 27 lncRNA signatures segregate the breast cancer subtypes in the Indian cohort

The PCA of the patients shows an immediate improvement over existing standard gene sets (PAM50, MammaPrint, and OncotypeDX) in the segregation of hormone receptor subtypes in the PCA. The clear separation of Hmod (moderate Her2 expression, ER/PR negative) from the other subtypes is noted, as is also visible in the heatmap of lncRNAs. These are of particular interest as Her2 specific lncRNAs (Figure 5g). Further, the triple-negative (TNBC) and triple positive (EPH) subtypes cluster surprisingly close together. The two Luminal A groups, ER and EP, do not cluster closely, indicating the heterogeneity observed within Luminal A tumours. When the combined list was checked for pre, and postmenopausal samples, the pattern observed for only the lncRNA list repeated as Hmod and ER were seen to be as a distinct cluster in postmenopausal samples, and an improvement from the previous signatures was observed in the premenopausal samples where EPH subtype segregated from other subtypes. (Supplementary Figure 3f) in the PCA. Having identified the lncRNAs specific to each subtype, to understand the functional significance of the lncRNA in breast cancer pathogenesis, we checked for the lncRNA-mRNA pairs that were co-expressed in all subtypes of breast cancer.

### Unique lncRNA-mRNA signature in Breast Cancer subtypes

To identify potential functions of the lncRNAs – identified potential cis-acting lncRNA-mRNA pairs based on the overlap on the chromosomes. Although LncRNA regulates gene expression in cis and trans, we focused on the lncRNA-mRNA pairs in cis and with an overlap of 1000bp. Hmod showed a maximum number of cis-acting lncRNA-mRNA pairs (809 downregulated) (909 upregulated), followed by ER (524down, 565 up), which was in contrast with mRNA expression alone. The gene -lncRNA pair found in the same orientation (5’-3’-5’-3’) vs. opposite orientation (5’-3’-3’-5’) is presented as a bar graph in (Figure 6a). To correlate the overall expression of mRNA and lncRNA in a subtype-specific manner, we performed a Pearson correlation. We found a minimal correlation in ER (r=0.15, p=8e-11), and other subtypes had no significant correlation. Interestingly, 91% correlation in the EPH subtype was observed when Pearson correlation analysis was performed using downregulated and upregulated cis lncRNA-mRNA pair separately. All other subtypes did not show a significant correlation.

**Figure 6:**
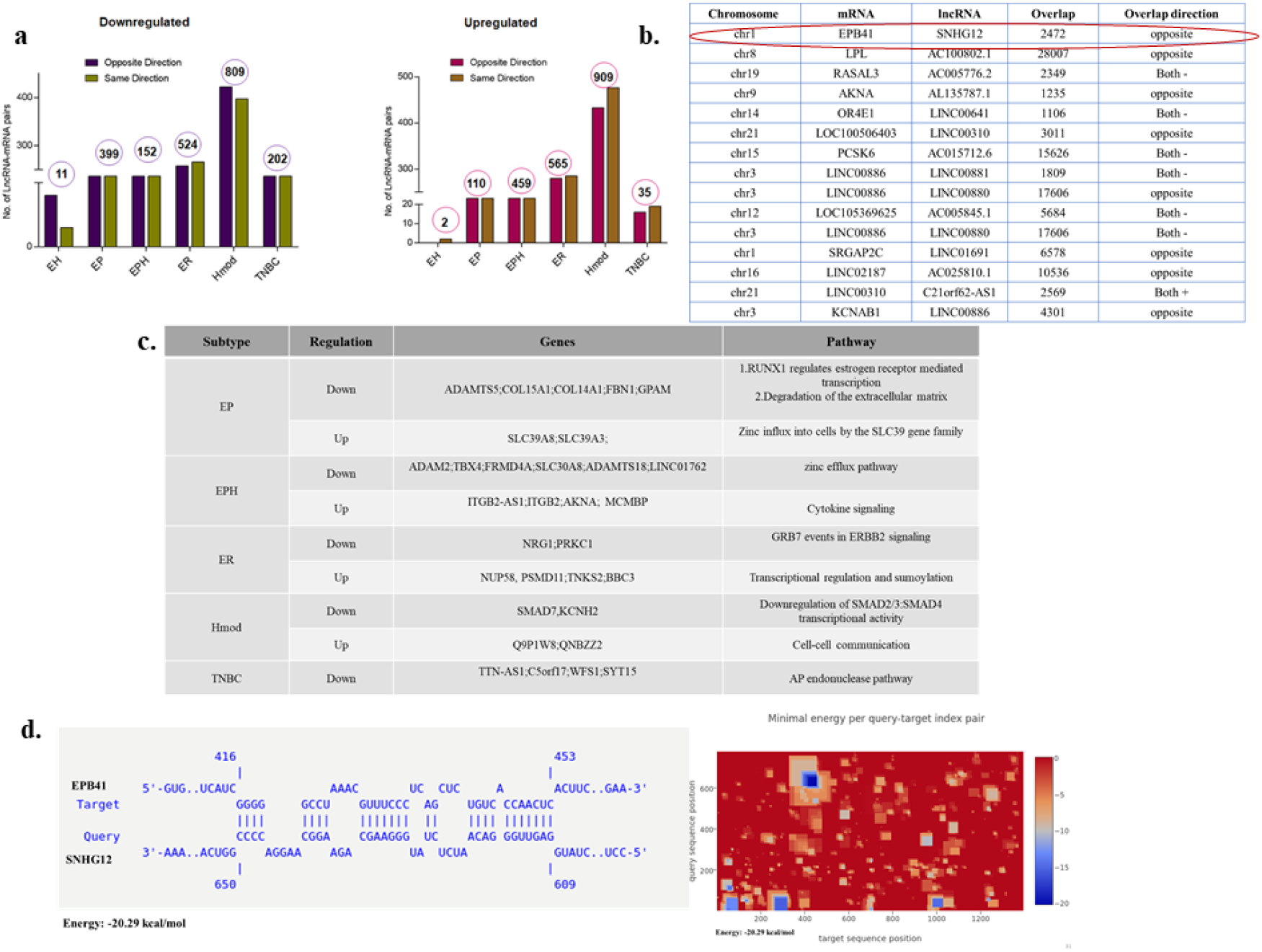
**a.** Bar graphs depicting No. of LncRNA-mRNA pairs in different subtypes of Indian Breast cancer patient samples. **b**. Table showing LncRNA and its corresponding mRNA pair obtained from different subtypes that were common to all the patients in the group. **c**. Table depicting subtype-specific pathways obtained from lncRNA-mRNA pairs **d**. A potential binding site between EPB41 (Target) and SNHG12 (Query) was identified by IntaRNA. Heatmap showing potential binding sites between EPB41 and SNHG12 in blue

Further subtype-specific lncRNA-mRNA pairs with Pearson correlation at least 90% were segregated (Figure 6b). The lncRNA was checked in the TANTRIC database for expression status and subtype specificity. Subtype-specific differences were observed in WAS-AS1, which is expressed highly in Basal in TCGA, whereas it is specific to Hmod and was not observed in Basal of the Indian cohort. SNHG12 is high in basal in TCGA datasets, whereas it is downregulated in EP in the Indian cohort. Linc00861 showed a downregulated expression pattern in both TCGA and the Indian cohort, whereas SLC39A8 was high in TCGA data and EP subtype in the Indian cohort and is associated with better survival.

The genes-lncRNA pair from each subtype were subjected to pathway analysis, and unique pathways were regulated in each subtype (Figure 6a). The downregulated pathways were zinc efflux transporters in EPH, whereas Zinc influx was upregulated in the EP subtype. Some of the underrepresented subtypes in mRNA were observed when the lncRNA-mRNA analysis was carried out. Combined analysis of lncRNA-mRNA returned some of the critical players in oncogenesis. To find out the lncRNA regulation of mRNA, several tools are available which can be used to identify the mode of action of lncRNA. We had noted that TSG SNHG12 was downregulated, and the cis gene ONC EPB41 was upregulated; we sought to narrow down on a mechanism using insilico methods.

### SNHG12 may regulate EPB41 specific to the EP subtype

To identify potential functions of the lncRNAs, potential cis-acting lncRNA-mRNA pairs were identified based on the overlap on the chromosomes. Among the lncRNAs, SNHG12 is known to be oncogenic and known to participate in proliferation, invasion, and metastasis in Breast Cancer tumours (40–42). In our cohort, SNHG12 was deregulated in the EP subtype. This lncRNA was picked up and looked for its mRNA pairs (Figure 6b). Erythrocyte Membrane Protein Band 4.1 (EPB41) was one of the interesting targets as it is known to play a role in invasion in other cancers (43–45). We wanted to see it’s binding and interaction with SNHG12.

INTERNA tool was used to check for the binding of the two. The results indicate a feasible binding between the lncRNA SNHG12 and the EPB41 (Figure 6c). While various regulatory functions of SNHG12 and EPB41 have been elucidated, the potential interaction between them remains unexplored and is a potential direction for further research. Similarly, to find out potential proteins that can bind to SNHG12 RNA, the eclip validated proteins were collated from the RNAct database (46) and checked for possible loss of oncogenic protein / PRC binding, which would block oncogene expression. We found 137 proteins that could bind to SNHG12 from the RNAct database out of which 5 of them (GTF2F1, APOBEC3C, DKC1, SUGP2, TIA1) were present in the gene list common to all EP patients.

### Validation of known cancer genes in Indian Breast cancer patients

We selected 5 Breast cancer-relevant genes, ALDH1A, BRCA, TP53, BCL2, and CD44, for validation using sybr green real-time PCR assays in n = 10 IDC samples. We observed that 40% of patients showed upregulation of ALDH1A and TP53. BCL2, an anti-apoptotic gene, was overexpressed in 50% of the patients. BRCA1 was commonly seen upregulated in 80% of the patients (Figure 7a). CD44 long and short-form levels were checked. It was observed that 60% of the patients showed upregulation of CD44l and s forms. Commonly deregulated lncRNA in Breast Cancer HOTAIR levels were also checked and it was observed that 77% of the patients showed a higher expression compared to their normal samples. When the patients were analysed for CD44l and s forms separately, it was seen that 50% of the patients had CD44l form high and CD44s form low, and 30% of the patients had CD44s form high and l form low (Figure 7b).

**Figure 7:**
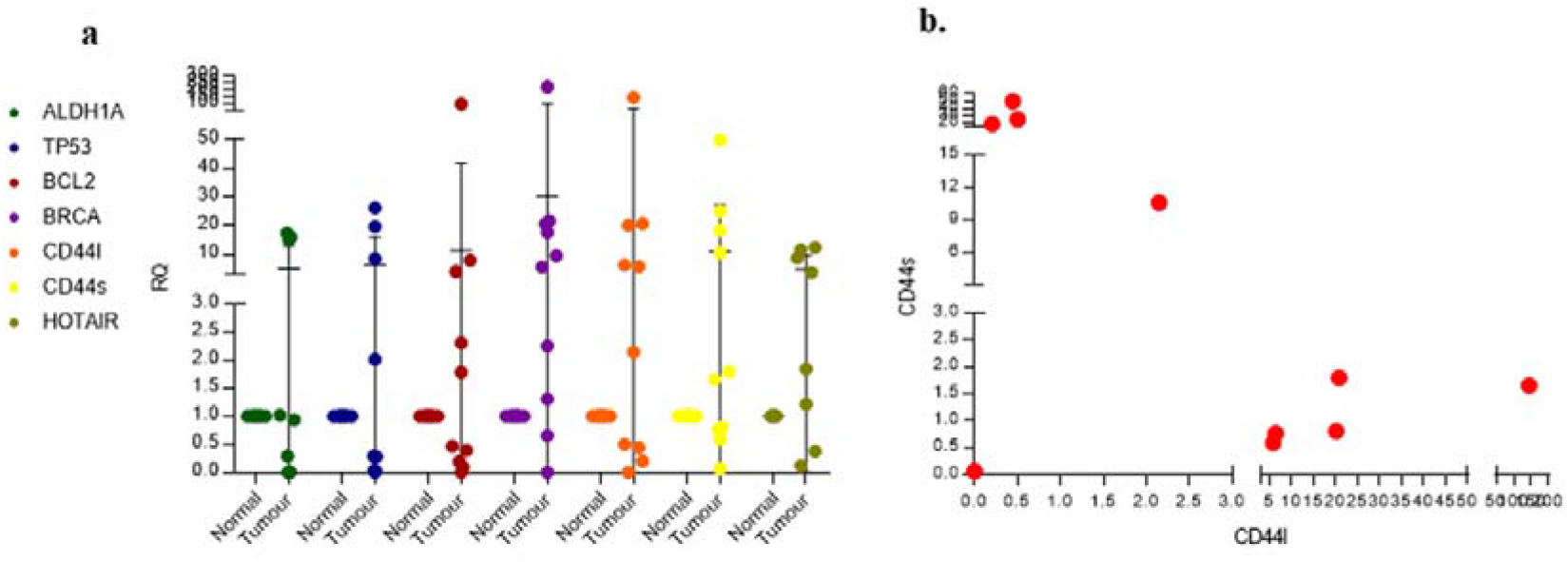
**a.** Real-time PCR dot plot depicting relative quantification for known Breast Cancer genes in Indian Breast Cancer patients. **b**. Scatter plot for checking co-expression of CD44 l and s forms

## Conclusion

Transcriptome sequencing and analysis of 17 Indian Breast cancer tumours and matched normal showed that already existing microarray gene signatures failed to segregate the samples into their subtypes using PCA. Every subtype showed a unique gene and pathway signature with minimum overlap. A unique set of differentially expressed onco and tumour suppressor lncRNA was identified for each subtype. Our data identified an mRNA-lncRNA gene set that could segregate pre and postmenopausal women with Breast Cancer. This is the first study reporting subtype-specific mRNA and lncRNA expression in Indian Breast Cancer patients. However, all these results need validation with a bigger sample size.

## Discussion

Breast cancer is a heterogenous and one of the major causes of death in women worldwide (7). Better insight into the molecular basis of the disease is possible when new approaches like Next-generation sequencing are used (18). Most of the Breast cancer data available in the repositories are from the Caucasian population (47). The gene signatures already available are from this population and population-specific changes are not very well addressed. Therefore, region-specific data generation with subtype information is necessary. One of the aims of our study was to generate breast cancer patient data for the Indian population categorized into 6 different subtypes based on hormone receptor status and to check for subtype-specific gene and lncRNA signatures. Our RNAseq data from 17 samples showed subtype-specific changes. The maximum alteration was observed in ER with 2572 genes downregulated and 1324 upregulated followed by were uniquely significantly downregulated and 1324 upregulated, followed by EP, Hmod, EPH, TNBC, and the last being EH. Among the deregulated pathways, ER-positive subtypes showed Keratinisation, RUNX3, AP2 family of genes regulating transcription, metabolism pathways. ER-negative tumours showed deregulation of ubiquitination, FGFR signaling, ECM interactions and notch signaling, collagen and cellular pathways. Very few gene expression studies have been reported from India to date. One of the very early studies by Thakkar AD et al., showed 108 genes differentially expressed in 31 ER-positive Breast tumours using microarray analysis (48). They found these genes were mostly involved in mRNA transcription and cellular differentiation pathways. Another study also used microarray technology and sequenced 29 tumours categorised into luminal, Basal and Her2 and 9 normal samples. They showed cell cycle, DNA replication, lipid metabolism PPAR signalling, focal adhesion and metastasis to be deregulated in Indian samples (49). Furthermore, pathways related to collagen, focal adhesion and ECM were reported to be deregulated in various cancers including breast tumours in other populations (50–55).

LncRNAs are a class of non-coding RNAs with lengths between 200 and 200,000 bases (19). They lack protein-coding features such as open-reading frames. They bear many similarities to mRNAs, often having multiple exons and undergoing post-transcriptional changes including splicing, polyadenylation, and 5′-capping (20). The dysregulation of lncRNAs has in several cases been found to be directly or indirectly associated with the hallmarks of cancers, mediated by other interacting partners including proteins, other non-coding RNAs, transcription factors and histone complexes (41). Studies done previously from the western population have shown HOTAIR lncRNA to be overexpressed in HER2+ breast cancers and HOTAIRM1 in basal□like breast cancers (56). LINC160 and DSCAM-AS1 were seen to be highly expressed in luminal A and B respectively (57,58). H19, MALAT, BC200, XIST and ATB, are the other lncRNAs frequently deregulated in breast cancer (59– 62). However, there is a dearth of explicitly Indian population-specific research evaluating lncRNAs in breast cancer. We analysed our sequenced data for lncRNAs and found uniqueness in differentially expressed lncRNA in different subtypes. ER subtype had the highest alterations in lncRNA followed by Hmod, EP and EH. TNBC and triple-positive (EPH) cancer showed comparable levels of differentially expressed lncRNAs. ATXN8OS, UCA1, SNHG12, SNHG5, LINC02487, TCL6, TET2-AS1, PTENP1−AS were some of the unique lncRNAs found in our cohort from different subtypes that were deregulated. A study from another Indian breast cancer showed ADAMTS9□AS2, EPB41L4A□AS1, WDFY3□AS2, RP11□295M3.4, RP11□161M6.2, RP11□490M8.1, CTB□92J24.3, and FAM83H□AS1 to be deregulated in early-stage Breast Cancer (22). Among the differentially expressed lncRNAs in our data, SNHG12 (small nucleolar host gene 12), a lncRNA present on chromosome 1 at the p35.3 region was looked into further. The length of SNHG12 is ∼1,867 bases coding for SNORA16A, SNORA61, SNORA66, and SNORD99 (63,64). SNHG12 has been implicated in various cancers, such as gastric, Triple Negative Breast Cancer, glioma, osteosarcoma. In Triple-Negative Breast Cancers, gastric, glioma, SNHG12 is high in expression (40,42,64–66). ER positive breast tumours in the TCGA data showed low expression of SNHG12 that correlated with our studies (67). This also indicates tumour and subtype-specific expression of SNHG12. In our data, SNHG12 was downregulated in the EP subtype hinting at its possible dual role both as an oncogene and tumour suppressor which needs to be further investigated. Through eclip data from the RNAct database (46), we found proteins that could bind to SNHG12 and among them, GTF2F1, APOBEC3C, DKC1, SUGP2, TIA1 genes were found in our list for EP subtype.

From the clinical analysis of our data, recurrent samples, grade 3 and stage 4 samples showed poor survival that correlated with the other population data. Her2 positive cancers showed poor survival in our data. A study from India with 3453 patients showed five-year overall survival to be 96.11% (95.12–97.1) in hormone receptor-positive/HER2 negative, 92.74% (90.73–94.8) in TNBC and 90.62% (88.17–93.15) in HER2 subgroups (68). However, in a study conducted by Pan et al., with Asian Breast tumours, Her2 positive cancers with an enriched immune score showed better survival (69). Low-grade HER2-positive breast cancer patients showed poor survival outcomes in European populations (70).

Our RNAseq data failed to segregate PCA PAM50, Oncodx and Mamaprint. However, when we separated pre and postmenopausal samples we could see minimum segregation. DNA microarray data from the Indian Breast cancers had shown segregation for PAM50 geneset in the study by Malvia S et al. Multiple Breast cancer patient RNAseq studies involving western populations has shown segregation for PAM50 gene set. A 25mRNA and 27-lncRNA gene set was derived from our data after iteratively performing segregation. There are multiple studies available from the western population having gene signatures for breast cancer (17,52,71–74). But none for the Indian population. The limitation of this study is the sample size. Nevertheless, it is the only study that shows an mRNA-lncRNA gene signature for the Indian population that is subtype-specific. This definitely shows some potential and a foundation for further studies. A larger sample size for sequencing and validation could be utilized next to strengthen the signatures obtained.

## Abbreviations

RNA: Ribonucleic acid
mRNA: messenger RNA
lncRNA: long noncoding RNA
BC: Breast Cancer
IDC: Invasive ductal carcinoma
TNBC: Triple-Negative Breast cancer
ER: Estrogen receptor
PR: Progesterone receptor
HER2: Human epidermal growth factor receptor
TCGA: The cancer genome atlas
PAM50: Prediction Analysis of Microarray 50
FFPE: Formalin-Fixed Paraffin-Embedded
GLOBOCAN: Global Cancer Observatory: CANCER TODAY
ROR: Risk of Recurrence Score
UCSC: University of California, Santa Cruz
SAM: Sequence alignment map
BAM: Binary alignment map
PCA: Principal Component Analysis
NCBI: National Center for Biotechnology Information
ONC: Oncogenes
TSG: Tumour suppressor genes
DEGs: Differentially expressed genes

## Acknowledgements

We thank Mainak Chatterjee for the help with validation work.

## Funding

Financial support was provided by The Department of Science and Technology Fund for Improvement of S&T Infrastructure in Higher Educational Institutions (grant no. SR/FST/LSI□5361/2012), The Department of Biotechnology, India, Glue grant (BTIPR23078/MED/29/1253/2017), and The Departments Information Technology, Biotechnology and Science and Technology, Government of Karnataka, India. MM is supported by The Senior Research Fellowship from the Department of Science and Technology□Innovation in Science Pursuit for Inspired Research, India (DST/INSPIRE Fellowship/2016/IF160535).

## Authors’ contributions

BC, MM, and SN designed the study and revised the manuscript. JST, MN provided patient samples, patient characteristics and disease outcome data. MM, SVG conducted the transcriptome analysis, AM performed clinical analysis and validation, BC, MM, SN interpreted the results, MM wrote the first draft of the manuscript, and prepared the figure and Tables. BC reviewed the results and their interpretation, and supervised the study. All authors reviewed the manuscript. All authors read and approved the final manuscript.

## Ethics approval and consent to participate

The study was conducted according to the guidelines approved by the Institutional Ethics Committee of the Institute of Bioinformatics and Applied Biotechnology. Informed consent was obtained from all subjects involved in the study.

## Competing interests

The authors declare that they have no competing interest

## Availability of data and material

Data will be available upon request

## Consent for publication

Not applicable

## Notes

### Competing Interest Statement

The authors have declared no competing interest.

