## Supplementary material for "A comprehensive study of mRNA and long noncoding RNAs in Indian Breast cancer patients using transcriptomics approach": Supplemntary Figure

| Sl No. | Sample No | Sample Type | Subtype | Total reads | File size (GB) |
| --- | --- | --- | --- | --- | --- |
| 1 | P1 | Normal | EPH | 27436900 | 2.1 |
| 2 | P2 | Tumour | EPH | 98457512 | 5.4 |
| 3 | P3 | Normal | EPH | 113058008 | 8.4 |
| 4 | P4 | Tumour | EPH | 41443626 | 3 |
| 5 | P5 | Normal | TNBC | 303036382 | 23.2 |
| 6 | P6 | Tumour | TNBC | 49464644 | 3.8 |
| 7 | P7 | Normal | EP | 177267404 | 13.4 |
| 8 | P8 | Tumour | EP | 53492744 | 4 |
| 9 | P9 | Normal | ER | 117412262 | 8.7 |
| 10 | P10 | Tumour | ER | 62919394 | 4.6 |
| 11 | P11 | Normal | ER | 142853800 | 9.6 |
| 12 | P12 | Tumour | ER | 76358772 | 5.8 |
| 13 | P13 | Normal | EH | 8466 | 0.066 |
| 14 | P14 | Tumour | EH | 68497432 | 5.1 |
| 15 | P15 | Normal | EH | 55460172 | 4.1 |
| 16 | P16 | Tumour | EH | 76834102 | 5.7 |
| 17 | P17 | Normal | TNBC | 39391050 | 3 |
| 18 | P18 | Tumour | TNBC | 2573894 | 0.2 |
| 19 | P21 | Normal | Hmod | 20074426 | 1.6 |
| 20 | P22 | Tumour | Hmod | 58847290 | 4.2 |
| 21 | P23 | Normal | EPH | 54953952 | 4 |
| 22 | P24 | Tumour | EPH | 25316472 | 1.9 |
| 23 | P25 | Normal | Hmod | 14950564 | 1.1 |
| 24 | P26 | Tumour | Hmod | 47365906 | 3.4 |
| 25 | P27 | Normal | TNBC | 81865694 | 5.9 |
| 26 | P28 | Tumour | TNBC | 74632110 | 5.4 |
| 27 | P29 | Normal | ER | 52675726 | 3.8 |
| 28 | P30 | Tumour | ER | 78357172 | 5.7 |
| 29 | P42T | Tumour | EP | 215220038 | 19 |
| 30 | P43N | Normal | EP | 78009304 | 6 |
| 31 | P43T | Tumour | EP | 25033980 | 2.1 |
| 32 | P44N | Normal | EPH | 54121062 | 4.8 |
| 33 | P44T | Tumour | EPH | 79673930 | 6 |

**Table 1:** Table showing sequencing details of all the samples. Total reads and file size for each sample is given.

**Supplementary Figure Legends**

**Figure 1:** Bubble plots showing significantly downregulated pathways in the six subtypes obtained from the Reactome database. Y-axis shows pathway terms and the x-axis is the pathway enrichment score. The size of the bubble represents the gene count of the pathway and the colour gradient of the bubble is based on the p-value

**Figure 2:** Bubble plots showing significantly upregulated pathways in the six subtypes obtained from the Reactome database. Y-axis shows pathway terms and the x-axis is the pathway enrichment score. The size of the bubble represents the gene count of the pathway and the colour gradient of the bubble is based on the p-value.

**Figure 3:** Principal Component Analysis of **a.** Pre- and post-menopause TCGA breast cancer patient samples with PAM50 **b.** Pre and post-menopause Indian breast cancer patient samples with Mamprint geneset. **c.** Pre and post-menopause TCGA breast cancer patient samples with Oncodx geneset. **d.** PCA plots showing segregation with selected mRNAs between pre- and post-menopause TCGA Breast Cancer patients **e.** PCA plots showing segregation with selected LncRNAs between pre and post-menopause Indian Breast cancer patients **f.** PCA plots showing segregation with selected mRNA-LncRNAs between pre and post-menopause Indian Breast cancer patients
